# Tissue-resident memory B cells augment local anti-cancer immunity via IgA

**DOI:** 10.64898/2026.08.08.743713

**Authors:** Abrar Samiea, Roxanne Bahn-Bales, Julia Vanderstreet, Maryam Al-Ghezi, Lina Gao, Malia Rettig, Zihan Guo, Rashi Yadav, Daniel O. Herzig, Sandy H. Fang, Liana Tsikitis, Adel Kardosh, Lauren B. Rodda, Ferdinando Pucci, Wesley Y. Yu, Rebekka Duhen, Joshua M. Moreau

## Abstract

Tissue-resident memory B cells (B_RM_) provide powerful localized protection against microbial infection in barrier tissues. It is unknown if analogous B_RM_ populations survey solid tumors and contribute to anti-cancer immunity. We profiled B cells from patients with colorectal cancer and cutaneous basal cell carcinoma and identified a CD69^+^ memory B cell population consistent with a tissue-resident phenotype. Integrative analysis of transcriptomic datasets identified an optimized signature enriched across cancer types. Tumor infiltrating B_RM_-like cells preferentially exhibited autoreactivity and their signature correlated with patient outcomes and response to immunotherapy. Skin and lung targeted vaccination established localized B_RM_ that provided IgA dependent organ specific protection upon tumor challenge in murine models. These findings establish B_RM_ as an active component of anti-cancer immunity via preferential reactivity to tumor associated self-antigens.

## Introduction

Non-lymphoid tissues harbor populations of lymphocytes that remain localized without recirculation and provide rapid site-specific protection, especially to support barrier immunity. These tissue-resident lymphocytes include memory T cells (T_RM_) that contribute to early tumor immunosurveillance and can mediate powerful anti-cancer immune responses(*1–3*). Analogous tissue resident memory B cells (B_RM_) have been identified within the lung following respiratory infection and vaccination(*4–11*). These cells rapidly differentiate into antibody-secreting cells upon antigen re-encounter and generate localized IgA-mediated recall responses that provide superior protection compared with systemic immunization(*5–8, 11*). Consistent with these specialized functions, lung B_RM_ are transcriptionally distinct from circulating and lymphoid memory B cells, expressing tissue-residency markers, including CD69, CCR6, and CXCR3, that partially overlap with those of T_RM_ cells(*5–8, 11*). Memory B cells expressing B_RM_-associated features, including CD69, have been identified across multiple human tissues, however definitive identification of B_RM_ in humans remains challenging due to the difficulty of directly tracking residency(*5, 12–14*).

Tumor-associated memory B cells and plasma cells are enriched across many human cancers and frequently localize within tertiary lymphoid structures, where they are generally associated with favorable clinical outcomes and improved responses to immune checkpoint blockade(*15–19*). Recent large-scale single-cell multi-omic and spatial profiling studies have revealed heterogeneity among tumor-associated B cells and identified class-switched memory B cell populations expressing tissue-associated molecules including FCRL4 and T-bet (*15, 16*). FCRL4, a member of the Fc receptor-like family previously linked to tissue B_RM_ cells (*20, 21*), has also been associated with favorable clinical outcomes in human cancers(*15*), suggesting that resident-like memory B cells analogous to infection-induced B_RM_ may exist within tumors. However, whether these populations represent true B_RM_ cells, how they are established within tumors, and whether they directly contribute to anti-tumor immunity remain unknown. Here, we combined analyses of transcriptomic data, patient derived B cells, and mouse models for tracking antigen specific cells to define the developmental requirements and functional role of B_RM_ in cancer. We identified a core B_RM_ transcriptional program enriched across multiple human cancers and demonstrated that skin and lung B_RM_ mediate site-restricted anti-cancer immunity via recognition of tumor expressed antigens.

### B_RM_ signature cells accumulate in human tumors and are enriched for autoreactivity

Despite characterization of lung B_RM_ cells during infection (*5–11*) and reports describing memory B cells within tumors(*15–17*), the presence and functional significance of B_RM_ within the tumor microenvironment has not been explored. To determine whether human tumors and adjacent tissues harbor B_RM_, we profiled B cells isolated from colorectal carcinoma (CRC) and cutaneous basal cell carcinoma (BCC) samples, along with patient matched adjacent tissues and peripheral blood mononuclear cells (PBMCs). Across most patients, CD69, a putative B_RM_ and T_RM_ marker associated with lymphocyte retention in tissues (*1, 5, 9, 14*), was enriched in B cells within tumors and matched adjacent tissues but largely absent from B cells in PBMCs (Fig. 1A). As CD69 can be transiently expressed following B cell activation (*22*), we next examined whether tumor-associated CD69^+^ B cells exhibited phenotypic characteristics previously associated with B_RM_ populations. Both human and murine B_RM_ cells are predominantly class-switched and preferentially express the tissue retention chemokine receptors CXCR3 and CCR6 (*6, 8–10*). Consistent with this phenotype, most CD69^+^CD19^+^ B cells within tumors and adjacent tissues displayed an antigen-experienced memory phenotype defined by the expression of CD27, loss of IgD, and low CD38. (Fig. 1B, S1A). These cells also exhibited reduced CD62L and were positive for both CCR6 and CXCR3 (Fig. 1B, S1A). To more comprehensively define tumor infiltrating B_RM_, we generated a cross-species B_RM_ transcriptional signature by identifying enriched genes in both experimentally established mouse lung B_RM_ cells (*8*) and human upper-airway B cells annotated as B_RM_ populations (*14*). This approach identified a conserved 80-gene B_RM_ signature, which included genes such as *FCRL5, FCRL4, CCR6, ITGAX, TBX21* and *ZEB2* that have previously been linked to B cell tissue residency (Fig. 1C, supplementary data file 1) (*4, 6, 8, 13, 14, 20, 21, 23, 24*). Interferon signaling was among the most enriched pathway, consistent with reports that IFN-γ is a driver of B_RM_ differentiation (Fig. 1D) (*25*). We next assessed the prevalence of our B_RM_ signature across human cancers using the multicancer single-cell B cell atlas generated by Ma *et al*. (*16*), comprising 9 cancer types where sufficient B cell data from both tumors and patient matched PBMCs was available. Significant enrichment of our B_RM_ score was evident in tumor infiltrating B cells across multiple malignancies (Fig. 1E).

**Fig. 1.**
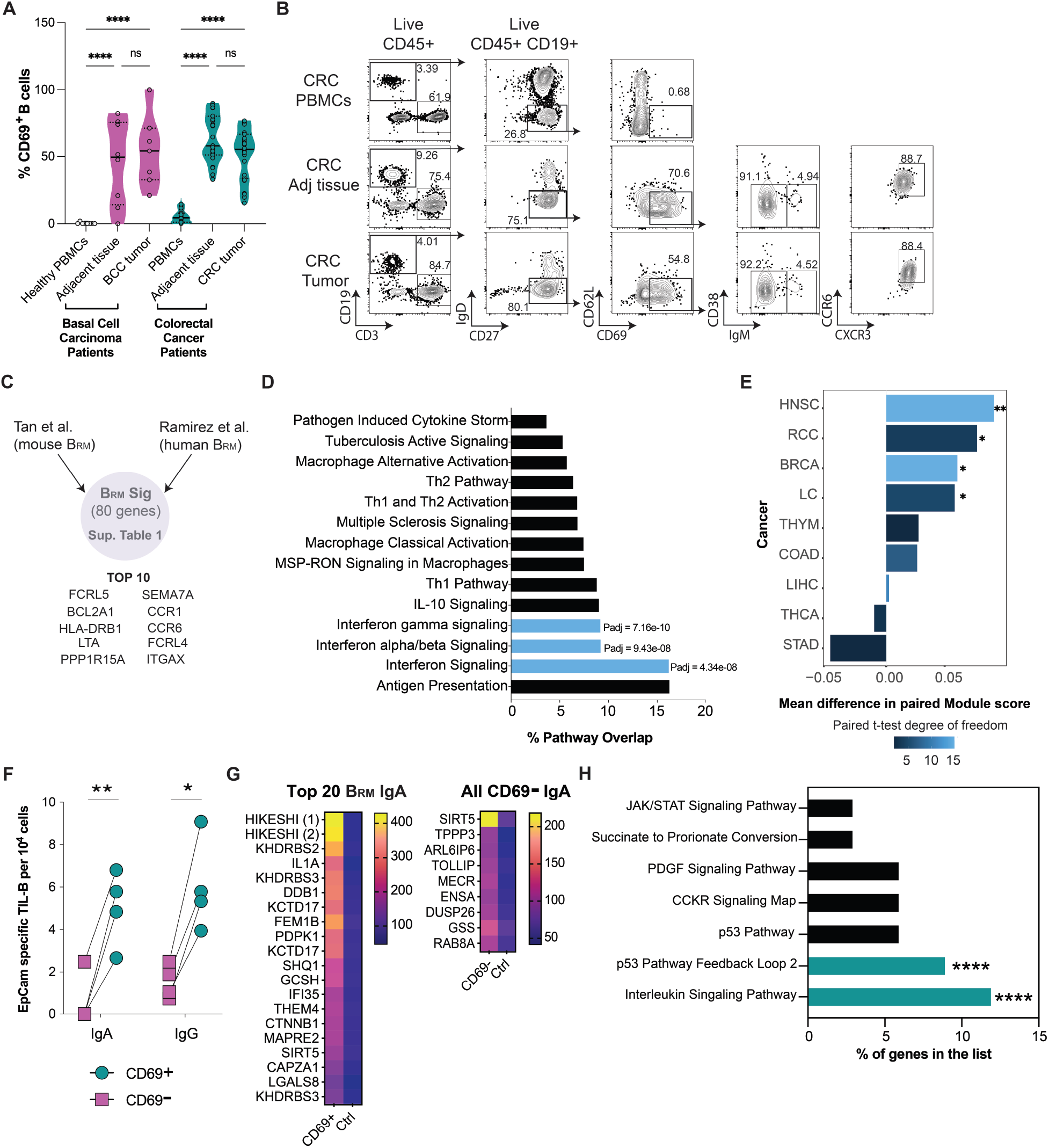
B_RM_ signature cells accumulate in human tumors and are enriched for autoreactivity. (**A**) Quantification of CD69^+^ B cell frequencies among live CD45^+^ CD19^+^ cells across CRC and BCC tumors, matched adjacent normal tissues, and matched or healthy PBMCs. Each point represents an individual patient sample (CRC: tumors, n = 24; matched adjacent tissues, n = 24; matched PBMCs, n = 24; BCC: tumors n = 7, matched adjacent tissues n = 7, healthy PBMCs n = 8). (**B**) Representative flow cytometry plots showing among live CD45^+^ cells from CRC tumors and patient matched tissues. (**C**) Top B_RM_ signature genes identified from mouse lung B_RM_ (*8*) and human upper airway (*14*) (**D**) QIAGEN Ingenuity Pathway Analysis of B_RM_ 80-gene signature. (**E**) Differential B_RM_ 80-gene signature module enrichment between tumor and peripheral blood across cancer types. Only cancer types with at least two tissue-matched patient samples were included in the analysis. (**F**) Antigen-specific antibody secretion by tumor-associated B_RM_ cells. Live CD45^+^ IgD^-^ CD69^+^ B_RM_ cells and matched CD45^+^ IgD^-^ CD69^-^ B cells were sorted from CRC specimens, differentiated in vitro, and analyzed by ELISpot. Quantification of EpCAM-specific IgA- and IgG-secreting cells per 10^6^ cultured cells. Each symbol represents an individual patient sample. Statistical significance was determined by paired two-tailed t-test. (**G**) Antigen specificity of antibodies produced by tumor-associated B_RM_ cells. Heatmaps showing the top antigen-specific IgA reactivities detected in culture supernatants from differentiated CD69^+^ IgD^-^ B_RM_ cells (left) and matched CD69^-^ IgD^-^ B cells (right) isolated from CRC specimens. Values represent normalized protein microarray signal intensities relative to control samples, with color intensity corresponding to the magnitude of antigen-specific IgA reactivity. (**H**) Pathway enrichment analysis of proteins preferentially recognized by B_RM_-derived IgA antibodies. Canonical pathway analysis was performed on proteins identified as targets of CD69^+^ B_RM_-derived IgA in the protein microarray dataset. Bar plots show the percentage of genes represented within each enriched pathway. Statistical significance was determined using pathway enrichment analysis. *P < 0.05, **P < 0.01, ****P < 0.0001.

B_RM_ cells can rapidly differentiate into antibody-secreting cells to generate local antibody responses (*5*). In several cancers intratumor B cells have been found to produce protective auto-antibodies specific for tumor associated antigens(*17, 19, 26–28*). We hypothesized that B_RM_ in human tumors would be enriched for autoreactivity. To test this, we isolated CD69^+^IgD^-^ cells and CD69^-^ IgD^-^ B cells from CRC tumor samples and measured their relative ability to produce antibodies against EpCAM, a CRC associated autoantigen (*29, 30*), by ELISPOT following activation. Putative B_RM_ cells generated significantly greater frequencies of EpCAM-reactive IgA- and IgG-secreting cells than matched CD69^-^ B cells (Fig. 1F). To more broadly assess the specificity of antibodies produced by tumor-associated B_RM_ cells, supernatants from differentiated CD69^+^ IgD^-^ B_RM_ cells and matched CD69^-^ IgD^-^ B cells isolated from CRC specimens were analyzed for IgG and IgA reactivity against 21,000 unique human proteins. Antibodies derived from CD69^+^ IgD^-^ cells exhibited broader autoantigen recognition, especially among the IgA isotype (Fig. 1G, supplementary data file 2). Pathway analysis of proteins recognized by CD69^+^ B cell derived IgA identified enrichment of p53-associated and interleukin signaling pathways (Fig. 1H). Notably, patients from whom these cells were isolated harbored a p53 tumor mutation (Supplementary data file 2), raising the possibility that B_RM_-derived antibodies contribute to local humoral responses directed against mutation-associated tumor antigens.

### B_RM_ augment local anti-tumor immunity

To dissect the functional role of B_RM_ cells in cancer, we developed a strategy to establish resident immune populations, reactive to distinct antigens, at separate tissue sites within the same mouse. Specifically, we used extracellular vesicles (EVs) derived from the murine squamous cell carcinoma MOC2 (*31*), engineered to express either transmembrane ovalbumin (mOVA), a model antigen recognized strongly by both B and T cells, or transmembrane hen egg lysozyme (mHEL), which in C57BL/6 mice is recognized exclusively by B cells (*32–34*). C57BL/6 mice were first primed intraperitoneally with mHEL-mOVA linked EVs before undergoing unilateral laser assisted epicutaneous vaccination with mHEL-mOVA EVs on one flank and mOVA EVs on the contralateral flank (Fig. 2A and B) (*35*). This approach selectively generated HEL-specific B_RM_ within defined skin regions while maintaining a shared systemic immune environment, thereby enabling direct comparison of local immune responses within the same animal. After a resting period of at least 35 days to allow B_RM_ establishment, mice were either analyzed at steady state or challenged bilaterally with mHEL-expressing MOC2 tumors. To track antigen-specific B cell responses, we used biotinylated purified recombinant OVA or HEL proteins tetramerized with fluorochrome-conjugated streptavidin (*36*), and quantified their distribution across spleen, skin, and tumor tissues (Fig. S2 A-D). Both OVA- and HEL-specific B cells were readily detected in vaccinated mice across all tissues examined. However, HEL-specific B cells were preferentially enriched within tumors arising at mHEL-mOVA vaccinated sites, whereas OVA-specific B cells were present at comparable frequencies across spleen, skin, and tumor tissues (Fig. S2 B-D). We next asked whether these HEL-specific B cells exhibited characteristics associated with B_RM_ populations. Consistent with this phenotype, HEL-specific B cells unlabeled by intravenously (IV) administered antibody were enriched at mHEL vaccinated sites and uniformly IgD^-^CD69^+^ (Fig. 2C, D). To determine whether enrichment of HEL-specific B_RM_ was site restricted, we assessed the presence of HEL- and OVA-specific IV^-^ B cell populations across mOVA and mHEL-mOVA vaccinated skin and tumors (Fig. 2D and S2E-G). HEL-specific IV^-^ B cells preferentially accumulated within mHEL-mOVA vaccinated skin and were further enriched in corresponding tumors while OVA-specific IV^-^ B cells were detected at comparable frequencies at both sites, consistent with our vaccination strategy (Fig. 2D and S2E-G), indicating localized establishment of HEL-specific B_RMs_.

**Fig. 2.**
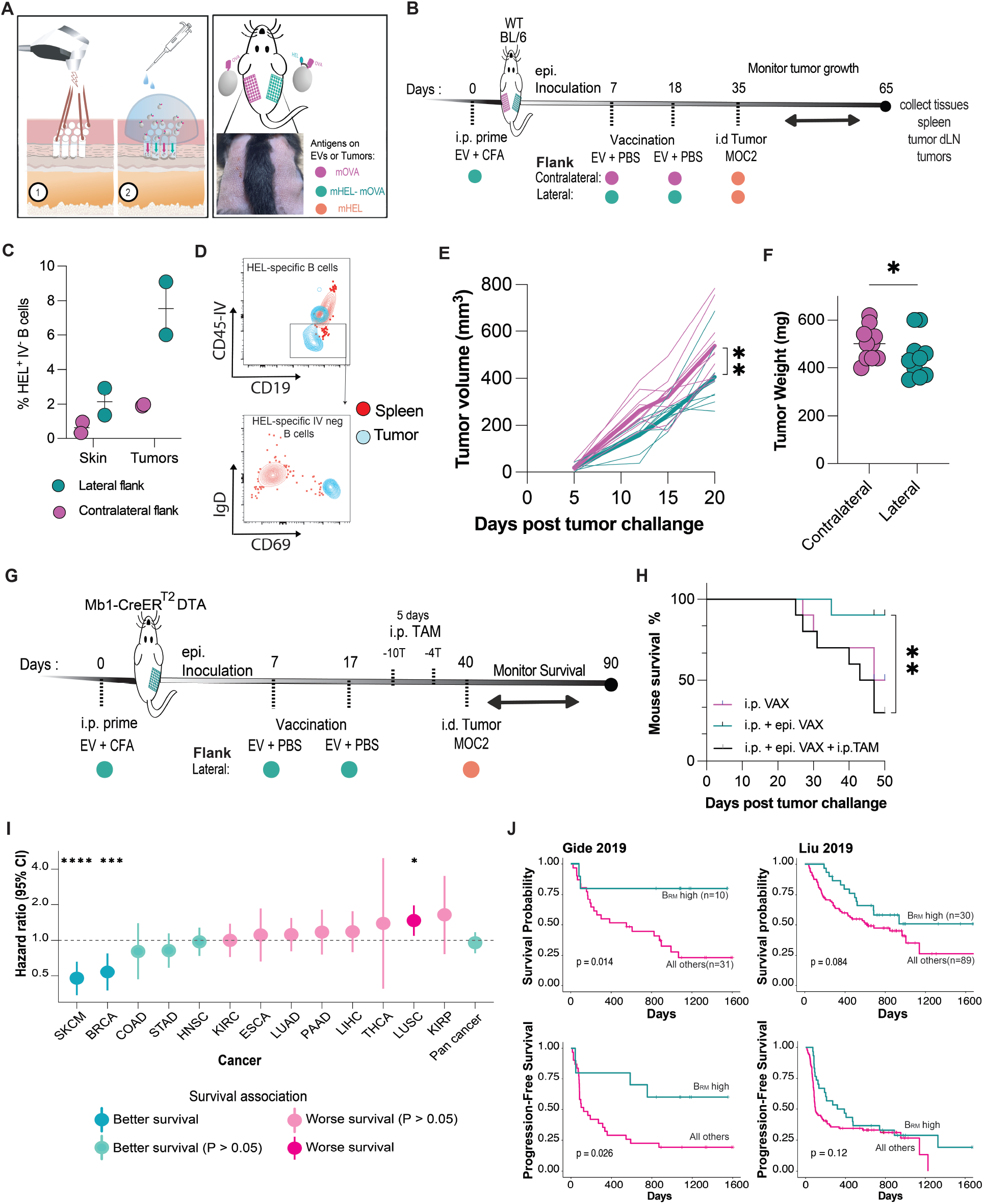
B_RM_ augment local anti-cancer immunity and correlate with patient outcomes. (**A** and **B**) Schematic overview of the laser assisted epicutaneous vaccination and tumor challenge strategy used to track B_RM_ responses. (**C**) Quantification of HEL-specific B cells in indicated tissues following vaccination. Each dot represents and independent experiment with 5 pool mice per group. (**D**) Representative flow cytometry plots showing CD69 expression on IV^-^ HEL-specific B cells (CD45^+^CD19^+^CD3^-^HEL^+^IV^-^) from vaccinated mice. (**E**) Tumor growth kinetics following bilateral intradermal implantation of MOC2-mHEL cells (n = 10 mice). Data are shown as individual tumors (dotted lines) and mean (solid lines) and are representative of three independent experiments. Growth curves were analyzed using two-way repeated-measures ANOVA with Geisser–Greenhouse correction. (**F**) Tumor weights at experimental endpoint. Each symbol represents an individual tumor and data are representative of three independent experiments. Endpoint tumor weights were analyzed using ratio paired *t*-test. (**G**) Schematic overview of B cell depletion strategy. (**H**) Kaplan-Meier survival analysis following B_RM_ establishment with or without B cell depletion across indicated treatment groups (i.p., n = 10; i.p. + epi., n = 10; i.p. + epi. + TAM, n = 10). Data are representative of one independent experiment. Survival curves were analyzed using a log-rank test. (**I**) Prognostic association of the B_RM_ signature across TCGA cancer cohorts. Hazard ratios of B_RM_-high compared to all others and 95% confidence intervals are shown for individual cancer types and pan-cancer analysis. *P* values of hazard ratios were derived from Cox proportional hazards models for each cancer type and from pan-cancer meta-analysis. (**J**) Kaplan–Meier plots of overall survival and progression-free survival in melanoma patient cohorts from Gide *et al*. (n = 41) (*41*) and Liu *et al*. (n = 121) (*42*), stratified based on B_RM_ 80-gene signature enrichment score. B_RM_ 80-gene signature enrichment was dichotomized into high (above the 75th percentile) versus all other samples. Survival curves were compared using two-sided log-rank tests. *P < 0.05, **P<0.01, ***P < 0.001, ****P < 0.0001.

Tumors implanted at sites enriched with HEL-specific B_RM_ cells exhibited significantly reduced growth and smaller endpoint weights compared to contralateral tumors within the same mice (Fig. 2E and F). To validate these findings via an independent approach, we employed an adoptive transfer model in which recipient mice received Hy10 B cells (*37*), engineered for reactivity to HEL, and OVA specific OT-II T cells (*38*), prior to unilateral mDEL-mOVA or mOVA EV vaccination. Following B_RM_ establishment animals recieved bilateral tumor challenge (Fig. S3A). DEL is a lower-affinity variant of HEL that binds the Hy10-B cell receptor with reduced affinity, enabling assessment of B_RM_ responses under more physiologic antigen-recognition conditions (*33*). Following bilateral challenge with mDEL-mOVA expressing tumors, Hy10-B cells preferentially accumulated within tumors arising at previously mDEL-mOVA vaccinated sites, whereas their abundance in the corresponding draining lymph nodes remained comparable between sites. In contrast, OT-II T-cell frequencies were similar between mDEL-mOVA and mOVA vaccinated tumors, as well as within their respective draining lymph nodes (Fig. S3B), indicating that local tumor control was not associated with altered T-cell accumulation. Tumors developing at previously mDEL-mOVA vaccinated sites exhibited reduced growth and endpoint tumor weights compared with contralateral tumors despite both sites being challenged with the same mDEL-mOVA expressing tumor cells (Fig. S3C and D). To directly determine if B cells mediated protection, we utilized Mb1-CreER^T2^; Rosa26-DTA mice (*39, 40*) to allow inducible B cell specific deletion after establishment of skin B_RM_ via epicutaneous vaccination (Fig. 2G). Administration of tamoxifen prior to MOC2 tumor challenge resulted in efficient depletion of B cells across tissues and completely ablated the skin B_RM_ population (Fig. S4). While localized epicutaneous vaccination improved survival compared to systemic vaccination this protection was lost following B cell depletion (Fig. 2H). Together, these findings provide evidence that B_RM_ provide immunosurveillance of local tumors and that they contribute to protective anti-cancer immunity.

### Tumor B_RM_-signature cells correlate with patient outcomes

To gain insight into the clinical relevance of B_RM_ cells, we leveraged TCGA datasets to evaluate the association between B_RM_ signature enrichment and patient 10-year overall survival across cancer types. Although heterogeneity was observed, B_RM_-associated transcriptional programs demonstrated strong positive correlations with outcomes in in cutaneous melanoma, breast invasive carcinoma, and colon adenocarcinoma (Fig 2I). Given the prominent association observed in melanoma, we next examined independent cohorts of melanoma patients treated with anti-PD-1 therapy (*41, 42*). In both the Gide *et al*. (n = 41) (*41*) and Liu *et al*. (n = 121) (*42*) cohorts, tumors enriched for the B_RM_ signature prior to immunotherapy were associated with improved overall survival and progression-free survival compared with all other tumors (Fig. 2J). These findings are consistent with our earlier observation that B_RM_-associated programs are enriched in tumor B cells across multiple human cancers (Fig. 1E) and with our experimental demonstration that antigen-specific skin B_RM_ cells promote local tumor control in mice (Fig. 2E-H).

### T cell help is required for establishment of protective skin B_RM_

Lung B_RM_ have been shown to depend on local antigen exposure and T cell CD40-CD40L mediated interactions during their establishment (*4, 10*). To determine if establishment of protective skin B_RM_ required T cell help, we leveraged the differential reactivity of T cells to HEL and OVA antigens in C57BL/6 mice (*32*). Animals were first primed intraperitoneally with mHEL-mOVA EVs to establish systemic HEL-specific memory responses and subsequently vaccinated with mOVA on one flank and mHEL on the contralateral flank (Fig. 3A). This approach allowed us to test whether local antigen recognition by B cells alone was sufficient to establish skin B_RM_ populations. Whereas OVA epitopes drive both B and T cell activation, in C57BL/6 mice, CD4^+^ T cell responses to HEL are severely limited preventing provision of T cell help (*32–34*). Antigen-specific B cell responses were quantified using OVA and HEL tetramers. OVA-specific B cells were readily detected in the spleen and tumors arising at mOVA-vaccinated sites. Conversely, HEL-specific B cells were nearly absent in flanks vaccinated with mHEL consistent with a requirement for T cell help (Fig. 3B-D). To determine whether these responses translated into anti-tumor immunity, mice were challenged bilaterally with HEL-expressing MOC2 tumors. Epicutaneous vaccination with mHEL alone was insufficient to provide protection (Fig. 3E and F). Although established B_RM_ populations can maintain protective humoral responses independently of continued T cell help (*4, 10, 11*), our findings suggest that like lung B_RM_ the establishment of B_RM_ in skin tissue requires local provision of T cell help.

**Fig. 3.**
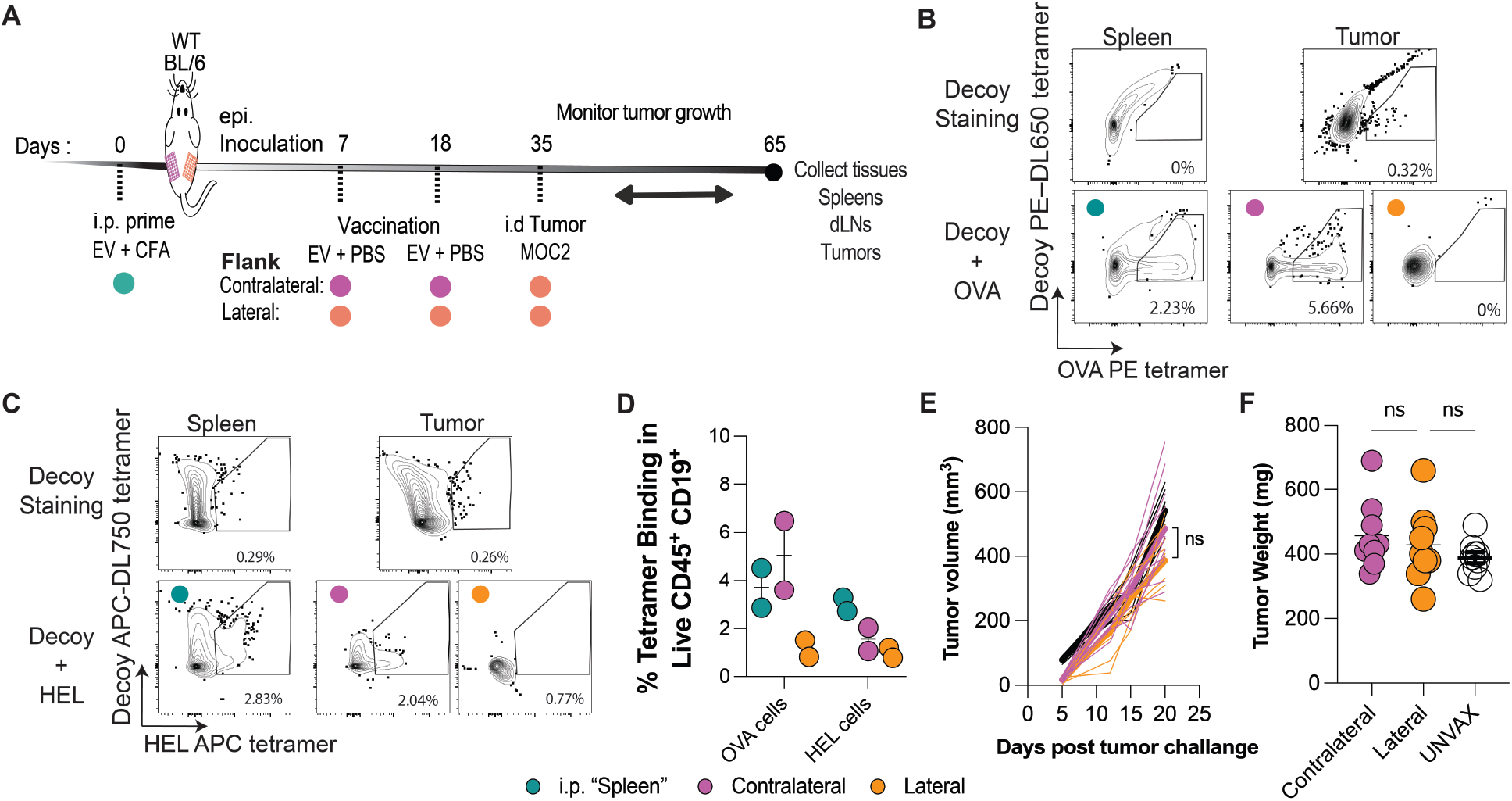
T cell help is required for establishment of protective B_RM_. (**A**) Experimental schematic for bilateral epicutaneous vaccination of mOVA and mHEL EVs. **(B** and **C)** Representative flow cytometry plots showing detection of OVA and HEL tetramer-specific B cells. Decoy tetramers were used to exclude nonspecific staining. (**D**) Quantification of tetramer-binding CD19^+^ B cells among live CD45^+^ cells in spleens and tumors. Results are pooled from two independent experiments. In each experiment, tissues from 10 mice were pooled, and each dot represents an independent experiment. (**E**) Tumor growth kinetics following bilateral intradermal implantation of MOC2-mHEL tumor cells into vaccinated lateral and contralateral flanks (n = 10 mice). Data are shown as individual tumors (dotted lines) or mean (solid lines) and are representative of two independent experiments. (**F**) Tumor weights at experimental endpoint. Each symbol is an individual tumor and data are presented are representative of two independent experiments. Paired tumor growth curves were analyzed using two-way repeated-measure**s** ANOVA with Geisser–Greenhouse correction. Endpoint tumor weights were analyzed using ratio paired *t*-test.

### B_RM_ derived IgA provides protection from lung metastasis

To determine if B_RM_ in barrier tissues other than the skin could confer protection against cancer challenge, we utilized an intranasal vaccination strategy to establish OVA specific B_RM_ in lungs. Mice were vaccinated with OVA by either the intranasal or intraperitoneal route and subsequently challenged with intravenous B16-OVA melanoma in the absence of T cells, enabling the selective assessment of B cell- and antibody-mediated protection (Fig. 4A). Consistent with previous studies demonstrating that intranasal, but not systemic, vaccination efficiently establishes lung-resident B_RM_, we readily identified class-switched, OVA specific parenchymal B cells following intranasal vaccination (Fig S5A and B) (*11*). Analysis of lungs following melanoma challenge revealed significantly reduced metastatic burden in animals receiving intranasal but not intraperitoneal vaccination (Fig. 4B-D). We next assessed if OVA specific antibody was detectible in lung tissue following vaccination. While anti-OVA IgA was readily detectible in the lungs of mice receiving intranasal vaccination, it was completely absent with systemic vaccination (Fig 4E). Conversely, anti-OVA IgG was present at high levels in the lungs of mice regardless of vaccination route (Fig. 4F). Given the selective induction of lung OVA specific IgA following intranasal vaccination, we asked whether IgA was required for B_RM_-mediated protection against melanoma metastasis. Vaccinated wild-type and IgA-deficient mice were challenged with intravenous B16-OVA melanoma in the absence of T-cell depletion. As expected, both naïve wild-type and IgA-deficient mice developed extensive pulmonary metastases following intravenous tumor challenge (Fig. 4G and H). While intranasal vaccination of wild-type mice markedly reduced metastatic burden, protection was significantly diminished in vaccinated IgA-deficient mice (Fig 4G and H). Although vaccination retained strong activity in the absence of IgA, this is consistent with parallel anti-OVA tissue resident memory T cell response (*43–45*). Concurrent depletion of CD4^+^ and CD8^+^ T cells, indicated that the protective effect of IgA was maintained in the absence of T-cell responses (Fig. S5C). Together, these findings demonstrate that local B_RM_-derived IgA contributes to protection against melanoma metastasis to the lung.

**Fig. 4.**
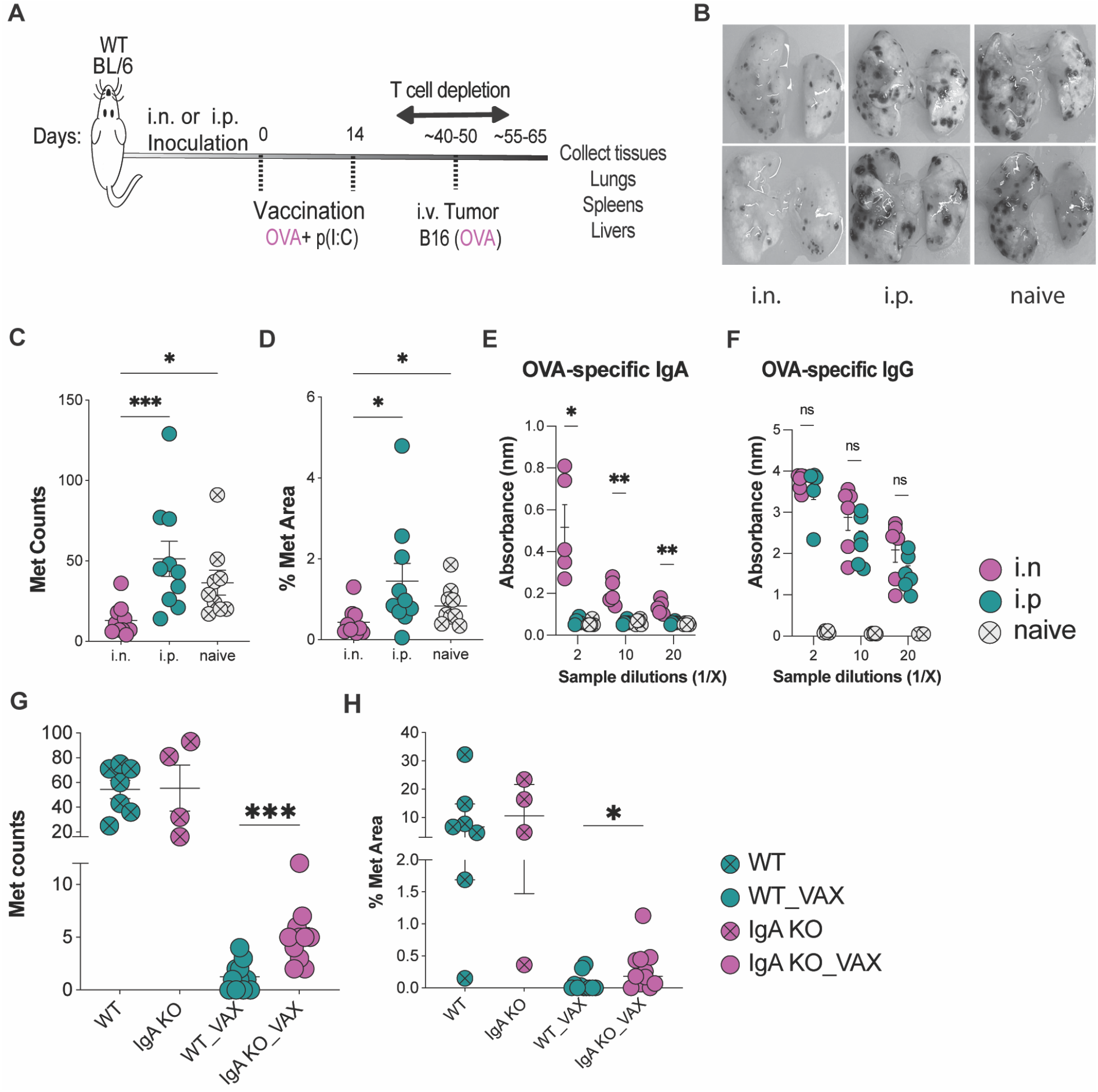
B_RM_ derived IgA provides protection from metastasis to the lung. (**A**) Schematic of lung vaccination and tumor challenge strategy. (**B**) Representative images of harvested lungs 14 days post-tumor challenge (naïve, n = 10; i.n., n = 10; i.p., n = 10). (**C** and **D**) Quantification of pulmonary metastatic burden of harvested lungs at endpoint based on the number of metastatic foci (**C**), and metastatic area (**D**). Each symbol corresponds to an individual lung and data are representative of two independent experiments. (**E** and **F**) The levels of OVA-specific IgA or IgG in the lung fluid following vaccination were measured by ELISA (n = 5 mice per group). Each symbol corresponds to an individual lung and data are representative of two independent experiments. (**G** and **H**) IgA-deficient and wild-type control mice were either left unvaccinated or vaccinated intranasally (i.n.) with OVA antigen combined with Poly (I:C) and boosted with same agents before tumor challenge. Lung metastatic burden was quantified the number of metastatic foci (**G**) and metastatic area (**H**). Each symbol corresponds to an individual lung and data are representative of one independent experiment. Metastatic burden analyses were performed using Kruskal-Wallis test with Dunn’s multiple-comparison testing. OVA-specific antibody ELISA data were analyzed using two-way ANOVA with Geisser–Greenhouse correction and Tukey’s multiple-comparison testing. Comparisons between vaccinated IgA-deficient and wild-type mice were analyzed using unpaired two-tailed Welch’s *t*-test. *P < 0.05, **P<0.01, ***P < 0.001

## Discussion

By combining profiling of B cells across multiple human cancers with a mouse vaccination model for establishment and tracking of antigen-specific memory lymphocytes in skin and lung, we identified an important role for localized B cell memory as a determinant of tumor control. We show that memory B cells with a tissue-residency transcriptional signature accumulate across cancers, preferentially exhibit autoreactivity, and correlate with patient outcomes, including responsiveness to immunotherapy in melanoma. Site-specific vaccination generating B_RM_ in skin and lung was sufficient to confer IgA-dependent, tissue-localized protection from tumor growth. These findings extend and unify prior previous research demonstrating the ability of B cells to acquire resident memory functionality as well as the protective potential of heterogeneous intratumoral B cells.

Tumor-reactive IgA and IgG ASCs have been described across multiple human cancers and are associated with favorable patient prognosis (*17, 26, 28, 46, 47*). While the origin of these local humoral responses has remained unclear, several lines of evidence implicate local B cell differentiation within tumoral associated TLS (*16, 18, 19*). Our findings suggest that B_RM_ may represent an important source of these effector cells, as CD69^+^ B_RM_ generated greater frequencies of tumor-reactive antibody secreting cells and exhibited broader autoreactivity than other tumor infiltrating B cells, consistent with an antigen-experienced population poised for rapid local recall responses. This is further substantiated by observations that ectopic tumor expression of autoantigens can drive intratumoral B cell affinity hypermutation of pre-existing autoreactive cells (*27*).

Previous studies of respiratory infection demonstrated that lung B_RM_ development requires local antigen encounter together with cognate CD4^+^ T-cell help mediated through CD40-CD40L interactions (*4, 10*). Additionally, T_fh_-derived IFNγ has been shown to promote B_RM_ differentiation by inducing T-bet expression and inducing CXCR3^+^ B_RM_ precursors (*25*). Our findings extend these developmental principles beyond the respiratory tract by demonstrating that the skin likewise supports the generation of B_RM_ in an antigen and T cell dependent manner. Skin HEL-specific B cells induced by site targeted vaccination with mHEL-mOVA localized within the tissue parenchyma and exhibited phenotypic features consistent with the non-recirculating B_RM_ population described in the lung, including expression of CD69. However, local antigen exposure alone was insufficient for B_RM_ establishment. Instead, productive B_RM_ responses required linked T cell activation via cutaneous mOVA vaccination, indicating that local helper T cell responses actively instruct tissue B_RM_ differentiation. This is consistent with recent reports that TLS can form in skin and support long-term B cell responses in this tissue (*48–50*). The conservation of these developmental requirements between the lung and skin suggests that common molecular pathways govern B_RM_ establishment across anatomically distinct barrier tissues.

Our study establishes tissue residency as an important organizing principle of anti-tumor B cell immunity and shows that this compartment can be deliberately programmed by tissue-directed vaccination. Given that current cancer vaccines predominantly target systemic immunity, strategies to establish or expand B_RM_ within barrier tissues represent an underexploited approach for durable, site-specific tumor control.

## Acknowledgments

The authors would like to thank members of the Oregon Health & Science University Immunology Group for thoughtful discussions and input. Data presented in this work was generated through support from the following OHSU Shared resources: Advanced Computing Center (RRID: SCR_009959), Flow Cytometry Shared Resource (RRID: SCR_009974), and the Knight Cancer Institute Cancer Data Science Shared Resource. We also thank the OHSU Department of Comparative Medicine. Results presented here are partly based upon data generated by the TCGA Research Network: https://www.cancer.gov/tcga.

## Funding

Elsa U. Pardee Foundation (JMM)

Collins Medical Trust (JMM)

Melanoma Research Alliance Young Investigator Grant 1253470 (JMM)

LEO Foundation LF-OC-23-001147 (JMM)

OHSU Knight Cancer Institute Faculty Startup Support (JMM)

NCI Cancer Center Support Grant P30 CA069533

## Author contributions

Conceptualization: AS, JMM

Methodology: AS, LG, FP, WY, RD, LBR, JMM

Investigation: AS, RB, JV, MA, MR, ZG, RY

Visualization: AS, RB, LG

Resources: FP, WY, DOH, SHF, LT, AK

Funding acquisition: JM Supervision: FP, RD, JMM

Writing – original draft: AS, JV, MA, JMM

## Data, code, and materials availability

All data are available in the main text or the supplementary materials. Transcriptomic datasets are available from cited publications. Code will be made publicly available.

## Supplementary Materials

Materials and Methods

Figs. S1 to S5

Table S1

References (*51–53*)

Data File S1 and S2

