## Supplemental Materials for "Tissue-resident memory B cells augment local anti-cancer immunity via IgA"

**The PDF file includes:**

Materials and Methods
Supplementary Text
Figs. S1 to S5
Table S1

### Materials and Methods

#### Clinical samples

Patients with CRC or BCC undergoing surgical resection were recruited through the Department of Dermatology and collaborating surgical oncology services at Oregon Health and Science University (OHSU) and were enrolled under OHSU Review Board-approved protocols (00025603 and 00022198). All participants provided written informed consent for tissue and blood collection. Fresh tumor and patient-matched adjacent tissue were collected at the time of surgery. For BCC specimens, fresh 6-mm punch biopsies of tumor and adjacent skin were analyzed. Peripheral blood was collected, when available, from CRC patients in EDTA tubes. In addition, healthy donor blood was collected. Specimens and PBMCs were processed into single-cell suspensions as described below and either used immediately or cryopreserved.

#### Analysis of single-cell RNA sequencing

Published processed single-cell RNA-seq datasets with cell-type annotations from Ma *et al. (16)* were downloaded at Cancer B cell blueprint (http://pancancer.cn/B/). Downstream analysis was performed in R using Seurat 5.3.0 *(51)*.

***$B_{RM}$ signature generation.*** To generate a conserved $B_{RM}$ transcriptional program from both human and mouse $B_{RM}$ signatures (**Fig. 1C**), $B_{RM}$ up-regulated genes identified in human upper airway single-cell RNA-seq data by Ramirez *et al. (14),* and the 200 top genes enriched in mouse lung $B_{RM}$ from Tan *et al. (8)* after conversion to corresponding human genes were examined. Genes identified in at least two independent datasets were retained, resulting in an 80-gene $B_{RM}$ signature comprising both genes conserved between human and mouse $B_{RM}$ and genes reproducibly identified across multiple human $B_{RM}$ datasets.

***$B_{RM}$ module score analysis across cancer types.*** To determine $B_{RM}$ signature enrichment across cancer types (**Fig. 1E**), $B_{RM}$ 80-gene signature module scores were analyzed using the published multicancer single-cell B cell atlas generated by Ma *et al. (16).* The integrated dataset comprised 474,718 B cells from 477 samples across 269 patients and donors spanning 20 cancer types, including head and neck squamous cell carcinoma (HNSC; n = 21), breast cancer (BRCA; n = 26), renal cell carcinoma (RCC; n = 6), lung cancer (LC; n = 29), colorectal adenocarcinoma (COAD; n = 78), hepatocellular carcinoma (LIHC; n = 15), thyroid carcinoma (THCA; n = 14), esophageal carcinoma (ESCA; n = 32), stomach adenocarcinoma (STAD; n = 7), ovarian cancer (OV; n = 4), pancreatic adenocarcinoma (PAAD; n = 3), cholangiocarcinoma (CHOL; n = 8), gallbladder carcinoma (GBC; n = 9), thymoma (THYM; n = 4), cervical squamous cell carcinoma and endocervical adenocarcinoma (CESC; n = 4), gastrointestinal stromal tumors (GIST; n = 2), neuroblastoma (NB; n = 3), bladder urothelial carcinoma (BLCA; n = 1), uterine corpus endometrial carcinoma (UCEC; n = 1), and cutaneous T cell lymphoma (CTCL; n = 2). Processed expression matrices and associated metadata, including tissue location, patient identifier, tumor type, treatment status, and sample origin, were obtained from the original publications where available. Cell clustering and annotations were retained as originally defined by Ma *et al. (16)*. Cancer types with fewer than two matched patient samples were excluded from downstream comparative analyses. The conserved $B_{RM}$ 80-gene signature module scores in each cell were calculated using "Seurat::AddModuleScore()" function with default parameters (genes in the signature set were scored relatively to 100 random control genes in the same expression bin across 24 bins).

For each cancer type, mean module scores were calculated separately for tumor-associated and peripheral blood B cells from matched patients, which were used as the unit of analysis. Mean paired differences were calculated as tumor minus blood module score, such that positive values indicated increased $B_{RM}$ signature enrichment in tumor-associated B cells relative to matched peripheral blood. Statistical significance within each cancer type was determined using paired two-tailed *t*-tests.

***Pathway enrichment analysis.*** Canonical pathway enrichment analysis of the $B_{RM}$ 80-gene signature was executed using QIAGEN Ingenuity Pathway Analysis (IPA). The $B_{RM}$ gene list was against the Ingenuity Knowledge Base to identify significantly enriched canonical pathways. Pathways were ranked by adjusted *P* value, and the top enriched pathways were displayed by percentage pathway overlap, defined as the proportion of genes in each pathway represented in the $B_{RM}$ signature.

***TCGA data analysis.*** The TCGA Toil re-computed expression data and patient metadata were obtained from the TCGA Pan-Cancer cohort through the UCSC Xena website (https://xenabrowser.net/). To model the predictive value of $B_{RM}$ on the ten-year overall survival of cancer patients (Fig. 2I), $B_{RM}$ 80-gene signature enrichment scores were calculated using the AUCell R package as previously described by Yang *et al. (15)*. Briefly, patients were stratified into $B_{RM}$-high and $B_{RM}$-low groups according to the median AUCell enrichment score within each cancer type (*52*), and then the total B cell abundance for each TCGA tumor sample was obtained. Cox proportional hazards models were then used to evaluate associations between $B_{RM}$ signature enrichment and overall survival, and hazard ratios with 95% confidence intervals were reported for each cancer type independently, where HR less than 1 indicated association with improved survival and HR more than 1 represented association with worse survival. In addition, Pan-cancer associations were subsequently estimated by random-effects meta-analysis. Kaplan–Meier survival curves were generated using the survival and survminer R packages.

***Immunotherapy datasets analysis.*** To explore of whether the enrichment of $B_{RM}$ signature was associated with immunotherapy response (**Fig. 2J**), we collected two bulk RNA-seq datasets of melanoma patients receiving anti-PD1 therapy. We kept only samples from tumors prior to immunotherapy. For the Gide *et al.* cohort *(40,41)*, pre-treatment anti-PD1 samples were retained, and ipilimumab plus anti-PD1 samples were excluded, resulting in 41 samples, including 19 responders. For the Liu *et al.* cohort *(41)*, 121 pre-treatment melanoma samples were analyzed. $B_{RM}$ 80-gene signature enrichment scores were calculated for each sample using AUCell. For survival analyses, samples were dichotomized into $B_{RM}$-high tumors, defined as those above the 75th percentile of $B_{RM}$ signature enrichment, and all other tumors.

### Mice

Mb1-CreER$^{T2}$ (JAX 033026), Rosa26 Ai14 (JAX 007914), Rosa26 DTA (JAX 009669), C57BL6/J (JAX 000664), and IgA-/- (Cyagen C001394) mice were purchased from Jackson laboratory or Cyagen biosciences and maintained in our laboratory. HEL-specific BCR-transgenic Hy10 mice (CD45.1 C57BL/6), maintained hemizygous for heavy or light chain transgenes, and TCR-transgenic OT-II mice (CD90.1 C57BL/6) were kindly provided by Dr. Ferdinando Pucci (OHSU) and used for adoptive transfer experiments. 6-16 week-old wild type 45.2-C57BL/6 mice, *Mb1-CreER$^{T2}$; Rosa-DTA* mice, or IgA KO female mice were used for tumor growth experiments. All mice were maintained in a specific pathogen-free environment, and all procedures were approved by the OHSU Animal Care and Use Committee.

### Tumor cell lines

Two murine tumor cell lines were used in this study: the oral epithelial squamous cell carcinoma cell line MOC2 engineered to express transmembrane model antigens (OVA, HEL, HEL-OVA, or DEL-OVA) *(31)*, and the melanoma cell line B16-MO4-OVA (Sigma, SCC420). MOC2 cells were cultured in E-medium at 37°C and 5% CO2 as previously described *(31)*. B16-MO4-OVA cells were cultured at 37 °C with 5% $CO_2$ in RPMI complete medium (RPMI-1640 supplemented with 10% FBS, 1× non-essential amino acids (GIBCO), 10 mM HEPES (GIBCO), 54 μM β-mercaptoethanol (Thermo Fisher Scientific), and 1 mg/ml Geneticin (G418; Sigma 345810)). For *in vivo* tumor challenge experiments, 5 X $10^5$ - 7 X $10^5$ MOC2 cells were injected intradermally (i.d.), whereas 5 X $10^5$ B16-MO4-OVA cells were injected intravenously (i.v.) to establish the experimental lung metastasis model.

**Extracellular vesicle (EVs) generation.** For the generation of $B_{RM}$ in the skin, purified transmembrane EVs were used for vaccination. EVs were purified from oral epithelial squamous cell carcinoma cell line MOC2 engineered to express transmembrane model antigens mOVA, mHEL, linked mHEL-mOVA or linked mDEL-mOVA as previously described *(31)*.

**Vaccination**

For intraperitoneal (i.p.) priming and induction of endogenous OVA- or HEL-specific immune cells, mice were immunized i.p. with 10 μg mHEL–mOVA EVs emulsified in complete Freund's adjuvant (CFA). For laser assisted epicutaneous vaccination, mice were anesthetized using inhaled 2% isoflurane in O2 during the microporation procedure, and dorsal flank hair was removed using depilatory cream at least 1 day before microporation. Skin microporation was performed using the P.L.E.A.S.E. laser device (Pantec Biosolutions) as previously described with modifications (*35*). Briefly, mice were positioned to expose the dorsal flank skin, and microporation was performed using the following settings: fluence, 11.9 J/cm²; pulse duration, 75 μs; repetition rate, 200 Hz; pulses per pore, 2; pore array size, 14 mm²; and pore density, 8%. Three microporated regions were generated per vaccination site. Following microporation, purified EVs expressing membrane-bound model antigens were diluted in sterile PBS without adjuvant and applied evenly onto the microporated skin surface. Mice received 15 μg EVs in 40 μL sterile PBS per flank distributed across the three microporated regions, with mHEL–mOVA, mDEL–mOVA, or mHEL EVs administered to one flank and mOVA EVs administered to the contralateral flank where indicated. The EV suspension was allowed to passively absorb into the microporated skin for approximately 10 min before mice were returned to their cages. A booster vaccination was administered 7 days after the initial vaccination, and tumor challenge was performed 14-35 days after vaccination as indicated in the experimental design. For intranasal vaccination, mice were anesthetized using inhaled 3% isoflurane with O2, and immunized as previously described with modifications *(11)*. Briefly, mice received intranasal administration of OVA antigen adjuvanted with Poly(I:C) in sterile PBS using a micropipette-based delivery method. Mice were positioned in a supine position at an approximate 60° incline, and the jaw was gently maintained in a closed position during inoculation to promote inhalation and prevent swallowing of the inoculum. Vaccines were delivered dropwise into alternating nares in a final volume of 40-60 μL. Mice were monitored until full recovery following immunization. In most experiments, mice received 50 μg OVA adjuvanted with 5–10 μg Poly(I:C), followed by a booster immunization 14 days later.

**In vivo B cells and T cell depletion**

To deplete CD4 and CD8 T cells, mice were treated intravenously with 300 μg each of anti-CD4 (GK1.5, BioXCell) and anti-CD8 (2.43, BioXCell) antibodies on days -4 and -1 before, and days +2 and +4 after tumor challenge. In addition, mice received 100 μg each of anti-CD4 and anti-CD8 antibodies intranasally on day -1 before challenge. T cell depletion efficiency was confirmed by flow cytometry where indicated. For inducible B cell depletion studies, Mb1-CreER$^{T2}$;Rosa26-DTA mice received intraperitoneal tamoxifen (MilliporeSigma) dissolved in corn oil at 75 mg/kg body weight once every 24 hours for 5 consecutive days.

**Adoptive transfer**
For adoptive transfer experiments, B and T cells were isolated from spleens and lymph nodes of HY10 mice (CD45.1$^{+/+}$) and OT-II mice (CD90.1$^{+/+}$) mice, respectively. Cells were processed into single cell suspensions. Cell quantity and purity were confirmed by flow cytometry following enrichment using negative selection kits (STEMCELL Technologies). Purified Hy10 B cells ($5 \times 10^5$) and OTII CD4 T cells ($2.5 \times 10^5$) were washed with PBS, resuspended in sterile PBS, and intravenously transferred into recipient Bl6.CD45.2 mice. One day after adoptive transfer, recipient mice underwent unilateral epicutaneous vaccination with mDEL-mOVA or mDEL EVs as described in the epicutaneous vaccination section.

**Tissue processing**
Mouse tissues were processed into single-cell suspensions as previously described *(53)*. To distinguish non-circulating tissue populations from circulating cells, mice received intravenous injection of 2 μg fluorochrome-conjugated anti-CD45 antibody 3 min before euthanasia, where indicated. Lungs were perfused by cardiac injection with 10 mL cold PBS before tissue collection. Skin and tumors were harvested, minced into smaller fragments in PBS, and centrifuged at 500 g for 5 min. Tissue supernatants were collected before digestion and stored at -80˚C for downstream protein analysis. Human tumors and matched adjacent tissue samples were processed similarly into single-cell suspensions. Under sterile conditions, specimens were cut into small pieces and digested in RPMI-1640 medium in the presence of hyaluronidase at 0.5 mg/ml (Sigma, H6254), collagenase at 1 mg/ml (Sigma, C5138), DNase at 30 U/ml (Roche, 04536282001) and human serum albumin (MP Biomedicals, IC08823051) at 1.5% final concentration. Cells were digested for 1 hr at 37°C under agitation. Cell suspensions were filtered through a 70 μm filter. Single-cell suspensions were cryopreserved until further analysis. Whole blood was collected into EDTA-coated tubes. Red blood cells were lysed using ACK lysis buffer for 3 min on ice, washed with cold PBS, and filtered through 40-μm strainers before downstream flow cytometric analysis. In some cases, human PBMCs were purified from whole blood over a Ficoll-Paque PLUS (GE Healthcare) gradient, washed and cryopreserved prior to analysis.

**Flow Cytometry**
For antigen-specific B cell identification, biotinylated recombinant OVA or HEL proteins were tetramerized with fluorochrome-conjugated streptavidin. Single-cell suspensions from mouse tissues were incubated with tetramers and decoy reagents, followed by magnetic column enrichment before starting surface staining as previously described *(36)*. Single-cell suspensions from mice or human samples were subsequently incubated with Zombie UV or Zombie Yellow Fixable Viability dye (Biolegend) for 15 min on ice to exclude dead cells, followed by Fc receptor blockade using anti-CD16/32 antibody (2.4G2) for 5 min at 4°C. Cells were then

stained with fluorophore-conjugated antibodies (**Table S1**) for 25 min at 4°C in the presence of Fc receptor blockade. Following surface staining, cells were fixed and permeabilized using the Foxp3/Transcription Factor Staining Buffer Set (ThermoFisher Scientific) according to the manufacturer's instructions. Stained cells were acquired using an Aurora flow cytometer (Cytek Biosciences).

**FluoroSpot**

For FluoroSpot assays, multiscreen 96-well plates (MAHAS4510, Millipore) were coated overnight at 4˚C with 15 µg of purified His-tag in PBS. Wells were washed and coated overnight at 4˚C with 10 µg of purified His-EpCAM antigen (SinoBiological, 10694-H08H-UE) in PBS. Plates were then washed with PBS and blocked with complete RPMI medium. Single-cell suspensions of human tissues and PBMCs were prepared as described above. Cells were thawed and $CD19^{+}$ $CD69^{+}$ $IgD^{-}$ and $CD69^{-}$ cells were sorted on a flow cytometer (BD Symphony S6 Sorter). Sorted cells were washed, diluted in complete RPMI medium, and cultured in CTL-Test B Medium containing resiquimod (R848), IL-2 (ImmunoSpot, CTL-mBPOLYS-200). Cells were incubated for 5 days at 5% 37˚C with 5% $CO_2$. 5 days post-incubation, cells and supernatants were collected. Supernatants were used for antibody reactivity array analysis (HuProt, v4.0, CDI LABS). Cells were then washed and plated onto EpCAM-coated FluoroSpot plates in complete RPMI medium and incubated overnight before plates were analyzed using the FluoroSpot human IgG or IgA Double-Color FluoroSpot kit according to manufacturer's instructions. Developed FluoroSpot plates were visualized and counted manually using the quickstart FluoroSpot guide package in Elispot CTL plate reader (Cellular Technology Ltd.).

**ELISA**

To detect HEL and OVA specific IgA or IgG, 96-well flat bottom microplates (Corning high binding) were coated overnight at 4˚C with recombinant HEL (10 µg/mL) or OVA (20 µg/mL) protein diluted in PBS. Plates were washed with PBS containing 0.05% Tween 20 and blocked for 1–2 h with PBS containing 5% FBS at room temperature. Tissue or tumor supernatant were followed by incubation for overnight at 4°C. After washing four times with PBS-Tween 20, bound antibodies were detected using horseradish peroxidase (HRP)-conjugated anti-mouse IgG or IgA (1:3000 dilution). After 1 hour incubation at room temperature, plates were washed and developed using 3,3',5,5'-tetramethylbenzidine (TMB) solution (eBioscience). Reactions were stopped with 1M hydrolic acid and absorbance were measured at 450 nm on a microplate reader.

**Statistics and software**

All flow cytometry plots and histograms were generated using FlowJo software (Version 10, Treestar). Statistical analyses were performed using GraphPad Prism (Version 11). The statistical significance of differences in mean values between groups were determined using unpaired or paired two-tailed Student's t test, or one-way or two-way ANOVA with appropriate correction or multiple-comparison testing, as specified in the figure legends. A *P*-value of less than 0.05 was considered significant.

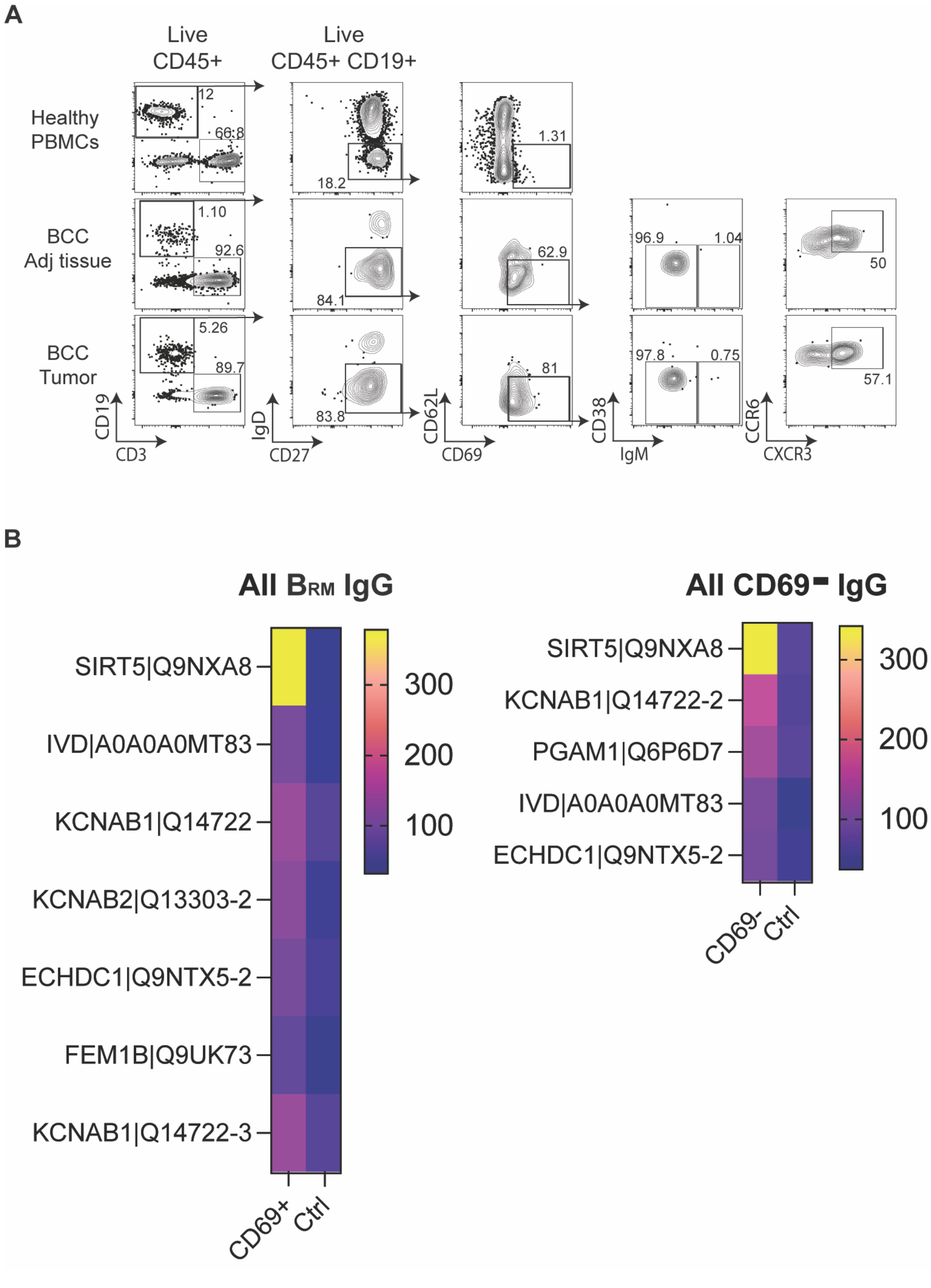


**Fig. S1. Enrichment of $B_{RM}$ cells in human tumors and adjacent tissues with autoreactivity.** (**A**) Representative flow cytometry plots showing memory and class-switched B cells ($IgD^-$ $CD27^+$) among live $CD45^+$ $CD19^+$ cells from BCC tumors, matched adjacent tissues, and healthy PBMCs. Memory B cells were further characterized for tissue-residency markers ($CD69^+$ $CD62L^-$), exclusion of germinal center and plasma cell phenotypes ($CD38^-$ $IgM^{-/+}$), and activation markers ($CCR6^+$ $CXCR3^+$). (**B**) Heatmaps showing the top antigen-specific IgG targets identified from culture supernatants of *in vitro*-differentiated live $CD45^+$ $IgD^-$ $CD69^+$ $B_{RM}$ (left) and $CD69^-$ B cells (right). Values represent normalized protein microarray signal intensities relative to control samples, with color intensity indicating the magnitude of antigen-specific IgG reactivity.

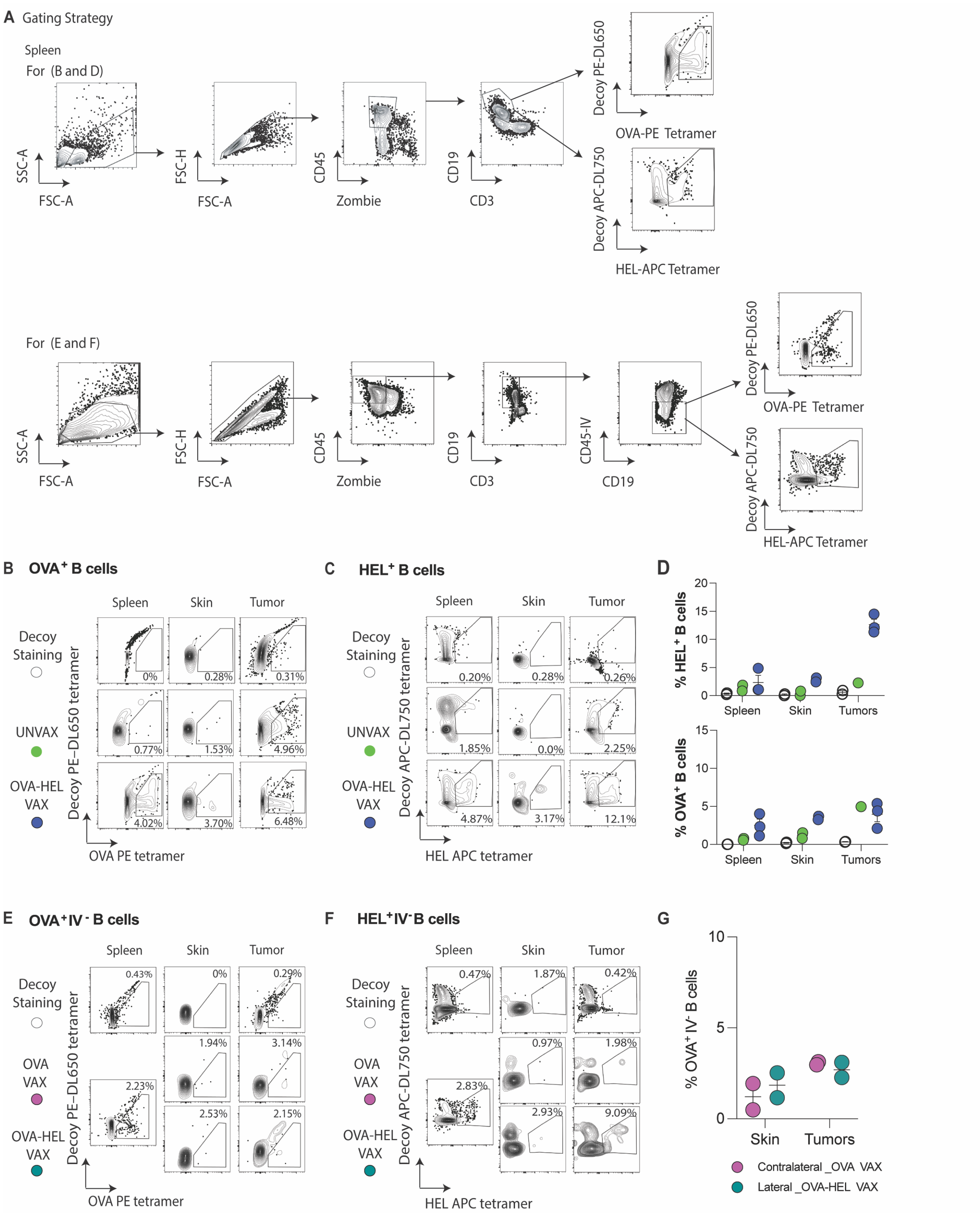


**Fig. S2. HEL or OVA antigen-specific B cell identification in different tissues following local B$_{RM}$ generation.** (**A**) Flow cytometry gating strategy with representative plots used to identify non-circulating HEL- or OVA- B cells from mouse spleen. Lineage exclusion included CD3ε staining followed by intravenous CD45 exclusion to identify non-circulating B cells.

Samples were acquired on a Cytek Aurora spectral flow cytometer. (**B** and **C**) Results obtained from gating in (**A**) to assess specific OVA or HEL B cells from spleen, skin and tumors of vaccinated C57BL/6 mice following the experimental strategy described in Fig. 1B. Skin tissues were collected 3 or 5 weeks after vaccination or mice were challenged intradermally with MOC2-mHEL tumor cells before collection of spleens and tumors to assess the presence of OVA (**B**) or HEL (**C**) specific B cells within vaccinated flanks. All harvested tissues were stained with decoy and tetramers. (**D**) Quantification of OVA-specific B cell frequencies is shown across indicated tissues. Results are pooled from two independent experiments. In each experiment, tissues from 5 mice were pooled, and each dot represents one pooled sample from an independent experiment.

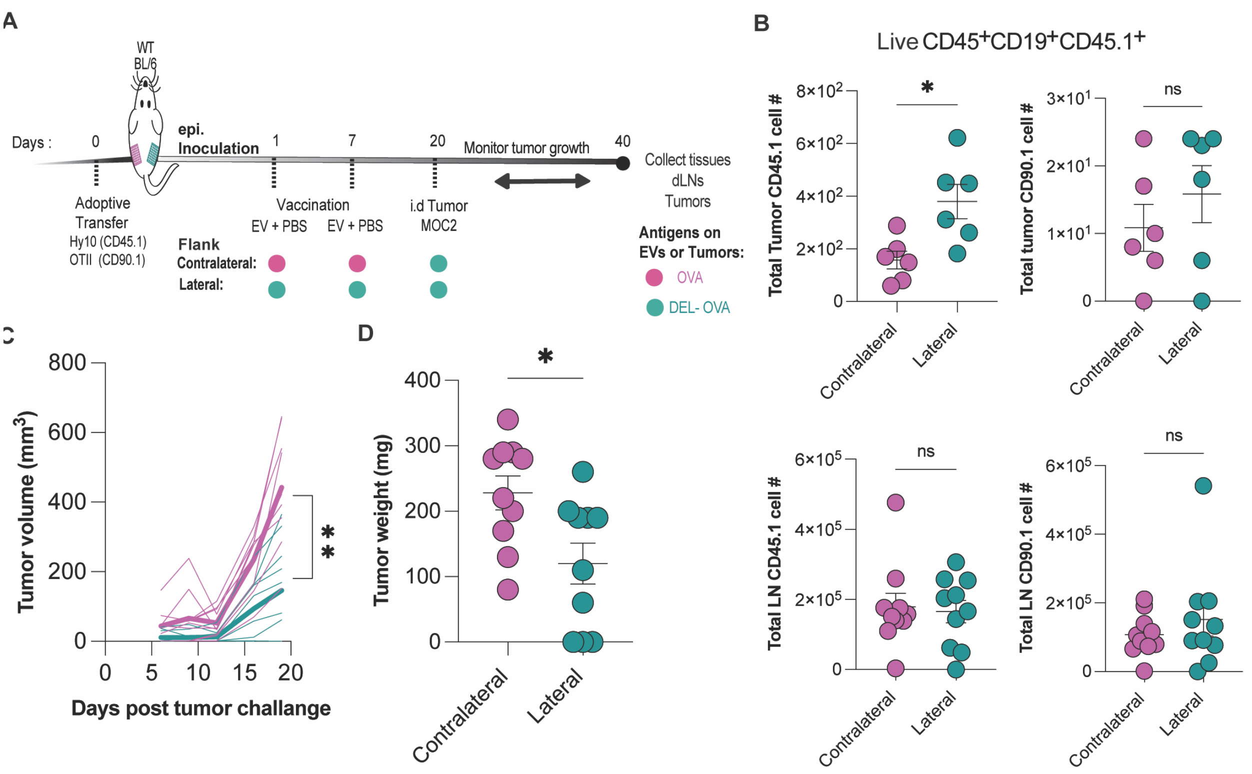


**Fig. S3. Local $B_{RM}$ established by Hy10 and OT-II cell adoptive transfer model augment anti-cancer immunity.** (**A**) Schematic overview of the adoptive transfer, EV vaccination, and tumor challenge strategy. Hy10 B cells and OTII T cells were adoptively transferred into C57BL/6 recipient mice, followed by unilateral administration of EVs expressing model antigens and boosted with same agents 10 days later, as indicated. Two to three weeks post-vaccination, mice were challenged intradermally on both vaccinated flanks with $5 \times 10^5$ MOC2 tumor cells expressing linked mHEL–mOVA antigens. (**B**) Quantification of total Hy10 and OTII B cells in tumors and draining lymph nodes are shown following vaccination and tumor challenge. (**C**) Tumor growth kinetics following tumor implantation in both flanks (n = 10 mice). Data are presented as individuals or mean. (**D**) Tumor weights were measured at experimental endpoint. Paired tumor growth curves were analyzed using two-way ANOVA with Geisser-Greenhouse correction. Total Hy10 and OT-II B cells and endpoint tumor weights were analyzed using paired two-tailed *t*-test. $P < 0.05$ was considered statistically significant. Data are representative of two independent experiments.

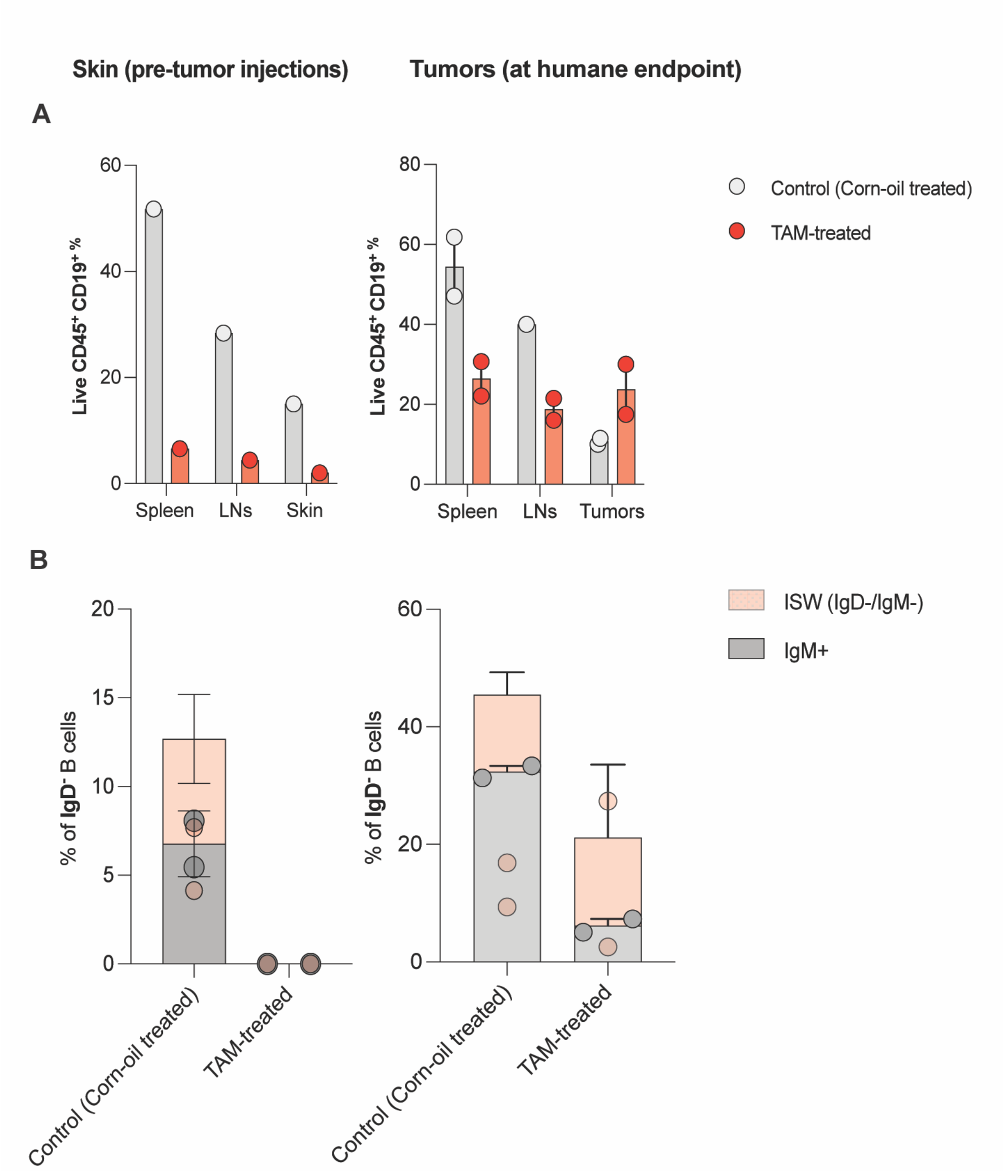


**Fig. S4. Demonstration of effective inducible B cell depletion following local B$_{RM}$ establishment.** (**A**) Quantification of $CD19^+$ B cell frequencies among live $CD45^+$ B cells in spleen, draining lymph nodes (dLN), and skin from mb1-CreER$^{T2}$;Rosa26-DTA mice treated with tamoxifen (TAM) or corn oil control following local HEL-OVA specific B$_{RM}$ establishment pre- and post- tumor challenge. (**B**) Quantification of class-switched ($IgD^-IgM^-$) and $IgM^+$ memory B cell populations in the skin and tumors of TAM-treated and control-treated mice (Skin; control n = 2 mice, in each indicated treatment group; 2 mice were pooled together); (Tumors; n= 2 mice in each indicated treatment group without pooling). Data are representative of one independent experiment.

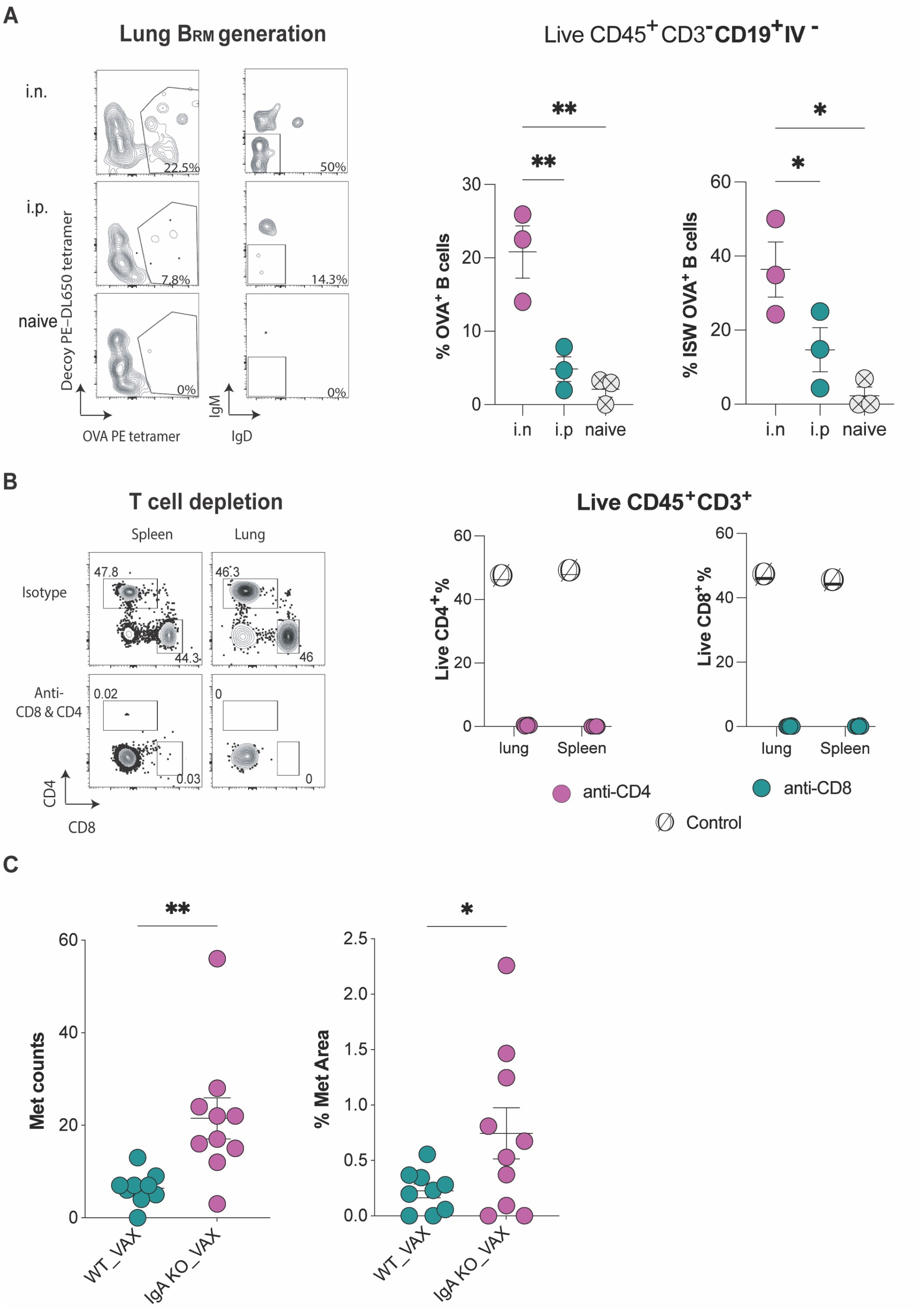


**Fig. S5. Demonstration of effective lung BRM establishment and T cell depletion. (A and B)** C57BL/6 mice were vaccinated intranasally (i.n.) or intraperitoneally (i.p.) with 50 µg OVA adjuvanted with 5 µg Poly(I:C) and boosted 14 days later with the same agents. **(A)** Representative flow plots and quantification of OVA-specific B cells and class-switched ($IgD^-$ $IgM^-$) OVA-specific B cells among non-circulating B cells from mouse lungs following vaccination. Gating strategy was adapted from Fig. S1A. Results are pooled from three independent experiments. In each experiment, lungs from two mice were pooled, and each dot represents one pooled sample from an independent experiment. (**B**) CD4 T and CD8 T cells depletion following intravenous and intranasal administration of anti-CD4 (GK1.5) and anti-CD8 (2.43) antibodies 25 days after vaccination. T cell depletion and quantification of CD4 T and CD8 T cells were assessed by flow cytometry. Data are representative of one independent experiment (isotype, n = 1; T cell depletion, n = 6). (**C**) IgA-deficient and wild-type control mice were either left unvaccinated or vaccinated intranasally (i.n.) with OVA antigen combined with and 5 µg Poly (I:C) and boosted with same agents before tumor challenge without T cell depletion (WT: VAX, n=9; $IgA^{-/-}$: VAX, n=10). Pulmonary metastatic burden was quantified by the number of metastatic foci and metastatic area. Data are presented as mean ± s.e.m. and are representative of one independent experiment. Comparisons between vaccinated IgA-deficient and wild-type mice were analyzed using unpaired one-tailed Welch's *t*-test.

### Table S1. Antibodies

| **Antibodies** | Source | Product ID |
|---|---|---|
| Flow cytometry: anti-mouse CD3 17A2 Spark Blue 574 | Biolegend | 100249 |
| Flow cytometry: anti-mouse CD45 30-F11 APC/Cyanine7 | Biolegend | 103115 |
| Flow cytometry: anti-mouse CD273 (PDL2) TY25 BUV563 | BD Biosciences | 741431 |
| Flow cytometry: anti-mouse CD138 281-2 PE/Fire640 | Biolegend | 142543 |
| Flow cytometry: anti-mouse IgM RMM-1 BV711 | Biolegend | 406539 |
| Flow cytometry: anti-mouse CD45R RA3-6B2 BV510 | BioLegend | 103248 |
| Flow cytometry: anti-mouse CD19 6D5 APC/Fire 810 | Biolegend | 115577 |
| Flow cytometry: anti-mouse CD80 MP5-20F3 Alexa Fluor® 488 | BD Biosciences | 561363 |
| Flow cytometry: anti-mouse CD73 TY/11.8 NovaFluor™ Yellow 730 | Thermo Fisher | M032T02Y07 |
| Flow cytometry: anti-mouse CD69 H1.2F3 Spark NIR 685 | Biolegend | 104557 |
| Flow cytometry: anti-mouse CXCR3 (CD183) CXCR3-173 NovaFluor™ Blue 610-70S | Thermo Fisher | M037T03B06 |
| Flow cytometry: anti-mouse CD279 (PD1) 29F.1A12 PE fire 810 | Biolegend | 135253 |
| Flow cytometry: anti-mouse IgD AMS 9.1 BV480 | BD Biosciences | 746452 |
| Flow cytometry: anti-mouse CD38 90 PerCP/Cyanine5.5 | Biolegend | 102722 |
| Flow cytometry: anti-mouse IgA IC10-1 BUV395 | BD Biosciences | 743299 |
| Flow cytometry: anti-mouse CD44 IM7 BUV661 | BD Biosciences | 741471 |
| Flow cytometry: anti-mouse CD43 S7 BUV737 | BD Biosciences | 612840 |
| Flow cytometry: anti-mouse CD62L MEL-14 BUV805 | BD Biosciences | 741924 |
| Flow cytometry: anti-mouse CD103 20000000 Alexa Fluor® 594 | Biolegend | 121428 |
| Flow cytometry: anti-mouse Ly6G 1A8 Spark Blue 550 | Biolegend | 127663 |
| Flow cytometry: anti-mouse TCR γ/δ GL3 BV605 | Biolegend | 118129 |
| Flow cytometry: anti-mouse CD3 17A2 BV750 | Biolegend | 100249 |
| Flow cytometry: anti-mouse CD8a RPA-T8 BV570 | Biolegend | 301037 |
| Flow cytometry: anti-mouse CD4 GK1.5 APC | Biolegend | 100411 |
| Flow cytometry: anti-mouse CD25 PC61 BV650 | Biolegend | 102038 |
| Flow cytometry: anti-mouse CD11c HL3 BUV737 | BD Biosciences | 612797 |
| Flow cytometry: anti-mouse CD11b M1/70 BV750 | BioLegend | 101267 |
| Flow cytometry: anti-mouse CD11a M17/4 BUV496 | BD Biosciences | 741071 |
| Flow cytometry: anti-mouse CD196 (CCR7) 29-2L17 BV785 | Biolegend | 129823 |
| Flow cytometry: anti-mouse CD5 53-7.3 PE-Cy5 | BioLegend | 100610 |
| Flow cytometry: anti-mouse CD45.1 A20 BV421 | BD Biosciences | 563983 |
| Flow cytometry: anti-mouse TIM1 RMT1-4 PE | BioLegend | 119506 |
| Flow cytometry: anti-mouse CD274 (B7-H1, PD-L1) 10F.9G2 PE/Dazzle™ 594 | BioLegend | 124324 |
| Flow cytometry: anti-mouse TCRb H57-597 Alexa Fluor® 700 | BioLegend | 109224 |
| Flow cytometry: anti-mouse CD19 6D5 APC-Fire 810 | BioLegend | 115578 |
| Flow cytometry: anti-mouse CD73 TY/11.8 PE-Dazzle 594 | BioLegend | 127234 |

| | | |
|---|---|---|
| Flow cytometry: anti-mouse CD69 H1.2F3 Spark NIR 685 | BioLegend | 104558 |
| Flow cytometry: anti-mouse CD183 (CXCR3) CXCR3-173 BV650 | BioLegend | 126531 |
| Flow cytometry: anti-mouse Ly-6G 1A8 Spark Blue 550 | Biolegend | 127664 |
| Flow cytometry: anti-mouse CD8a 53-6.7 BV570 | BioLegend | 100740 |
| Flow cytometry: anti-mouse CD4 RM4-4 BV605 | BioLegend | 116027 |
| Flow cytometry: anti-mouse CD80 16-10A1 BV421 | BD Biosciences | 562611 |
| Flow cytometry: anti-mouse CD138 281-2(RUO) BB515 | BD Biosciences | 566207 |
| Flow cytometry: anti-mouse CD43 S7 BV750 | BioLegend | 747277 |
| Flow cytometry: anti-mouse CD45 30-F11 PerCP-Cy5.5 | BioLegend | 103131 |
| Flow cytometry: anti-mouse GL7 GL-7 (GL7) eFluor 450 | Thermo Fischer | 48-5902-82 |
| Flow cytometry: anti-mouse CD38 90 PE-Fire 700 | BioLegend | 102747 |
| Flow cytometry: anti-mouse IgG GOT214A BUV615 | BD Biosciences | 752758 |
| Flow cytometry: anti-mouse TCR b H57-597 BV510 | BioLegend | 109234 |
| Flow cytometry: anti-mouse CD4 Ultra Violet GK1.5 | BD Biosciences | 613006 |
| Flow cytometry: anti-mouse CD80 Violet 16-10A1 | BioLegend | 104725 |
| Flow cytometry: anti-mouse IgG Violet Poly4053 | BioLegend | 405327 |
| Flow cytometry: anti-mouse CD3 Blue 17A2 | BioLegend | 100276 |
| Flow cytometry: anti-mouse ki67 Yellow-Green 11F6 | Biolegend | 151218 |
| Flow cytometry: anti-mouse FoxP3 Blue MF-14 | Biolegend | 126405 |
| Flow cytometry: anti-mouse CD45R RA3-6B2 AF700 | Biolegend | 103231 |
| Flow cytometry: anti-mouse IgD 11-26c.2a BV510 | Biolegend | 405723 |
| Flow cytometry: anti-mouse F40/80 BM8 BV785 | Biolegend | 123141 |
| Flow cytometry: anti-mouse CD45.2 104 (RUO) PerCP/Cyanine5.5 | BD Biosciences | 552950 |
| Flow cytometry: anti-mouse CD45R (B220) AF 700 RA3-6B2 | BioLegend | 103232 |
| IV Labeling for Flow cytometry: anti-mouse CD45 I3/2.3 FITC | Biolegend | 147710 |
| IV Labeling for Flow cytometry: anti-mouse CD45 30-F11 (RUO) APC R700 | BD Biosciences | 565478 |
| Invivo T Cell Depletion: anti-mouse CD4 GK1.5 | BioXCell | #BE0003-1 |
| Invivo T Cell Depletion: anti-mouse CD8 2.43 | BioXCell | #BE0061 |
| Flow cytometry: anti-humanCD45 BUV395 HI30 | BD Biosciences | 563791 |
| Flow cytometry: anti-humanViability Zombie UV [N/A] | Biolegend | 423108 |
| Flow cytometry: anti-humanCD45RA BUV563 HI100 | BD Biosciences | 612926 |
| Flow cytometry: anti-humanIgM BUV661 UCH-B1 | BD Biosciences | 750365 |
| Flow cytometry: anti-humanCD69 BUV737 FN50 | BD Biosciences | 564439 |
| Flow cytometry: anti-humanCD80 BUV805 L307.4 | BD Biosciences | 742046 |
| Flow cytometry: anti-humanCD279 (PD-1) BV421 NAT105 | Biolegend | 367422 |
| Flow cytometry: anti-humanCD196 (CCR6) Pacific Blue G034E3 | Biolegend | 353439 |
| Flow cytometry: anti-humanCD11c BV480 B-ly6 | BD Biosciences | 566135 |
| Flow cytometry: anti-humanCD62L BV510 DREG-56 | Biolegend | 304844 |
| Flow cytometry: anti-humanCD8 BV570 RPA-T8 | Biolegend | 301038 |

| | | |
|---|---|---|
| Flow cytometry: anti-humanIgD BV605 IA6-2 | Biolegend | 348232 |
| Flow cytometry: anti-humanCD183 (CXCR3) BV650 G025H7 | Biolegend | 353730 |
| Flow cytometry: anti-humanCD24 BV711 ML5 | Biolegend | 311136 |
| Flow cytometry: anti-humanCD185 (CXCR5) BV750 J252D4 | Biolegend | 356942 |
| Flow cytometry: anti-humanCD138 BV786 MI15 | BD Biosciences | 743501 |
| Flow cytometry: anti-humanT-bet Alexa Fluor 488 525831 | R&D | FAB53851G |
| Flow cytometry: anti-humanCD3 Spark Blue 550 SK7 | Biolegend | 344852 |
| Flow cytometry: anti-humanCD27 NovaFluor Blue 610-70S O323 | ThermoFisher | H012T03B06-A |
| Flow cytometry: anti-humanCD4 PerCp RPA-T4 | Biolegend | 300528 |
| Flow cytometry: anti-humanCD273 (PDL2) PerCP-Cy5.5 MIH18 | BD Biosciences | 564256 |
| Flow cytometry: anti-humanCD307d (FcRL4) PE 413D12 | Biolegend | 340204 |
| Flow cytometry: anti-humanCD197 (CCR7) PE-Dazzle 594 G043H7 | Biolegend | 353236 |
| Flow cytometry: anti-humanCD19 PE-Cy5 HIB19 | Biolegend | 302209 |
| Flow cytometry: anti-humanCD103 PE/Fire 700 Ber-ACT8 | Biolegend | 350240 |
| Flow cytometry: anti-humanCD25 PE-Cy7 BC96 | Biolegend | 302612 |
| Flow cytometry: anti-humanCD307e (FcRL5) APC 509f6 | Biolegend | 340306 |
| Flow cytometry: anti-human Foxp3 Alexa Fluor 700 PCH101 | ThermoFisher | 56-4776-41 |
| Flow cytometry: anti-humanCD38 APC/Fire 750 HB-7 | Biolegend | 356626 |
| Flow cytometry: anti-humanCD73 NovaFluor Red 755 AD2 | ThermoFisher | H016T03R06-A |
| FluoroSpot: anti-human CD45 BUV395 HI30 | BD Biosciences | 563791 |
| FluoroSpot: anti-human CD8 BUV805 SK1 | BD Biosciences | 564913 |
| FluoroSpot: anti-human CD3 BV510 OKT3 | Biolegend | 317331 |
| FluoroSpot: anti-human IgD BV605 IA6-2 | Biolegend | 348231 |
| FluoroSpot: anti-human CD4 AF488 RPA-T4 | Biolegend | 300519 |
| FluoroSpot: anti-human CD19 PE-Cy5 HIB19 | Biolegend | 302209 |
| FluoroSpot: anti-human CD27 APC M-T271 | Biolegend | 356409 |
| FluoroSpot: anti-human CD45 BV711 HI30 | BioLegend | 304050 |
| FluoroSpot: anti-human CD19 PE HIB19 | Biolegend | 302208 |
| FluoroSpot: anti-human CD69 APC-R700 FN50 | BD Biosciences | 565154 |
| Cell sorting: anti-human CD45 HI30 BV711 | BioLegend | 304050 |
| Cell sorting: anti-human CD19 HIB19 APC-Fire 810 | BioLegend | 302271 |
| Cell sorting: anti-human CD69 FN50 Spark NIR 685 | BioLegend | 310957 |
| Cell sorting: anti-human IgD IA6-2 BV480 | BD Horizon | 566138 |
| Cell sorting: anti-human CD3 OKT3 eFluor 450 | Invitrogen | 48-0037-42 |
